# UV-excited autofluorescence uncovers multicopy suppressors of a Cdc14-associated meiotic defect

**DOI:** 10.64898/2026.11.14.766682

**Authors:** George B. Garside, Carolina Pascual-Silva, Isabel Acosta, Jesús A. Carballo

## Abstract

The Cdc14 phosphatase plays essential roles in meiotic chromosome metabolism and cell-cycle progression, yet the cellular pathways capable of compensating for reduced Cdc14 function remain poorly understood. To explore these connections, we performed a multicopy suppressor screen in *Saccharomyces cerevisiae*, using a mutant *CDC14* allele that severely impairs sporulation. UV-excited ascospore-wall autofluorescence enabled the rapid screening of approximately 14,400 transformants, while subsequent microscopy provided quantitative validation of sporulation phenotypes. Photographic image processing facilitated candidate recovery, and a complementary computational analysis of archived images established the reproducibility of fluorescence-associated detections across different acquisition conditions. Genetic screening and independent plasmid retransformation identified four genomic-library plasmids that reproducibly enhanced sporulation. Molecular characterisation of the strongest suppressors revealed genomic regions associated with chromatin regulation and nuclear-pore organisation. Further analysis prioritised two particularly promising candidate genes: *RCO1*, encoding a component of the Rpd3S histone-deacetylase complex, and *POM34*, encoding an integral membrane nucleoporin. Suppressor activity was retained in a reduced *POM34*-containing construct, strengthening its candidacy for further functional investigation. Together, these findings establish an effective fluorescence-guided strategy for multicopy suppressor discovery and reveal promising genetic connections between Cdc14-dependent meiotic progression, chromatin regulation and nuclear architecture.

## Introduction

In *Saccharomyces cerevisiae*, the completion of meiosis depends on the successful coordination of chromosome metabolism with two successive nuclear divisions and the differentiation of four haploid spores. Following a single round of DNA replication, homologous chromosomes pair, synapse and undergo recombination during an extended prophase I (Zickler and Kleckner 1999; Marston and Amon 2004). The connections established between homologues must then be resolved in time for their segregation at the first division, while sister chromatids separate during meiosis II (Alonso-Ramos and Carballo, 2024). Formation of a mature ascus requires more than the execution of these chromosome movements: spore membranes must assemble around the haploid products and develop a specialised wall that protects them outside the vegetative cell cycle (Briza et al., 1990a; Neiman, 2011). Meiotic development also entails coordinated changes in gene expression (Chu et al., 1998; Chu and Herskowitz, 1998), alongside changes in chromatin organisation and nuclear architecture, including remodelling of nuclear pore complexes (Neiman, 2011; King et al., 2019, 2023). Sporulation efficiency provides a convenient measure of meiotic progression and spore differentiation (Neiman, 2011), including defects that are not readily detected by colony growth.

The *CDC14* gene was identified in classical cell-division-cycle mutant studies (Culotti and Hartwell, 1971). The Cdc14 phosphatase occupies a central position in the regulatory network that coordinates cell-cycle transitions in budding yeast. Much of our understanding of Cdc14 has emerged from mitotic studies, in which its timely availability is regulated by nucleolar sequestration and release (Visintin et al., 1999; Shou et al., 1999; Stegmeier and Amon, 2004; Alonso-Ramos and Carballo, 2024). For much of the cell cycle, Cdc14 is retained in the nucleolus through association with Net1/Cfi1, restricting its access to substrates elsewhere in the cell. Its mobilisation during anaphase involves the FEAR network and mitotic exit network (MEN), whose sequential and partially distinct contributions help couple substrate dephosphorylation to chromosome segregation, spindle disassembly and mitotic exit (D’Amours and Amon, 2004; Rock and Amon, 2009). The spatial and temporal regulation of substrate access is therefore essential for Cdc14 function (Visintin et al., 1999; Shou et al., 1999; Alonso-Ramos and Carballo, 2024). In mitotic cells, Cdc5 has been implicated in Net1 phosphorylation and FEAR-network interactions, while its later degradation favours the return of Cdc14 to the nucleolus (Shou et al., 2002; Rahal and Amon, 2008; Visintin et al., 2008).

During meiosis, Cdc14 functions in cells that reorganise chromosome structures after an extended prophase and undergo two successive divisions without intervening DNA replication. Early genetic studies implicated Cdc14 and components of the FEAR pathway in meiotic spindle disassembly and nucleolar division, demonstrating that mechanisms first characterised during mitotic exit also contribute to the physical progression of meiosis (Marston et al., 2003; Buonomo et al., 2003). Other studies examined MEN regulation during meiosis (Attner and Amon, 2012) and established a Cdc14-independent requirement for Cdc15 in spore morphogenesis (Pablo-Hernando et al., 2007). Chromosome recombination, the distinctive behaviour of homologues and the requirements of the second nuclear division create additional opportunities for Cdc14-dependent regulation, including events that precede the most conspicuous redistribution of the phosphatase during anaphase I. The timing of Cdc14 mobilisation is particularly relevant at the transition from meiotic prophase to chromosome segregation, when recombination intermediates must be resolved and chromosome-associated structures are remodelled (Alonso-Ramos et al., 2021).

DNA double-strand breaks introduced by Spo11 initiate interactions between homologous chromosomes (Keeney et al., 1997; Lam and Keeney, 2015), and their repair generates intermediates that must be processed into appropriate crossover and non-crossover products (Hunter, 2015). The resolution of joint molecules is coordinated with cell-cycle progression, partly through phosphorylation-dependent control of structure-selective nucleases (Matos et al., 2011). In our previous work, Cdc14 contributed to the activation and nuclear localisation of the Yen1 endonuclease, the resolution of recombination intermediates and crossover formation, particularly in backgrounds where alternative processing activities were compromised. These findings implicate Cdc14 in meiotic chromosome metabolism as well as division-associated events (Alonso-Ramos et al., 2021). Additional evidence that nucleolar Cdc14 organisation responds to meiotic recombination context has reinforced the biological connection between chromosome events and nuclear architecture (Rodríguez-Jiménez et al., 2026).

In our previous work, we characterised a C-terminally triple-HA-tagged allele of *CDC14*, referred to here as *cdc14-HA3* (designated *cdc14-HA* in Alonso-Ramos et al., 2021). Although the mutant retains substantial mitotic function, homozygous diploids exhibit a pronounced sporulation defect. During meiosis, Cdc14-HA3 protein abundance is reduced, and the mutant displays defects in recombination-intermediate resolution and chromosome segregation. These findings link the sporulation phenotype to impaired meiotic chromosome metabolism, rather than establishing an isolated defect in spore morphogenesis. Importantly, increasing the copy number of the tagged allele itself partially restored sporulation (Alonso-Ramos et al., 2021).

These observations prompted us to investigate whether increased dosage of other genomic regions could alleviate the *cdc14-HA3* sporulation defect. We chose a multicopy genomic-library suppressor screen as one genetic approach to this question, allowing us to survey regions beyond established Cdc14 regulators.

A large genomic-library screen required a rapid means of identifying potentially sporulating transformants. The yeast ascospore wall contains dityrosine, which exhibits characteristic fluorescence under UV excitation and consequently offers a means of recognising colonies that produce spores without immediately examining every transformant microscopically (Briza et al., 1986; Briza et al., 1990b). Dityrosine fluorescence has also been used to monitor spore formation in studies of meiotic checkpoint regulation (Acosta et al., 2011). We therefore combined UV inspection with photographic image processing and microscopic scoring of recovered colonies.

Here we present a multicopy genomic-library screen in the homozygous *cdc14-HA3* background, integrating fluorescence-guided recovery with microscopic quantification, genetic assessment and independent plasmid retransformation. We establish the photographic and image-processing procedures used to identify potentially sporulating colonies and evaluate a computational approach that measures the reproducibility of fluorescence-associated detections across screening plates. We then examine the phenotypes of selected isolates, demonstrate plasmid-associated suppression for a reproducible subset and characterise the genomic fragments associated with the strongest responses.

## Results

### UV autofluorescence provides an effective strategy for identifying sporulating transformants

To identify multicopy suppressors of the *cdc14-HA3* sporulation phenotype, we transformed the homozygous mutant diploid GGY8 with the YEp13 genomic library. The sporulation-proficient heterozygous *CDC14*/*cdc14-HA3* diploid GGY10 served as a reference throughout the screening procedure. Transformants were selected on SC-Leu medium and replica-plated onto KAc-Leu for sporulation. Candidate colonies were identified by exploiting the characteristic UV-excited autofluorescence associated with dityrosine accumulation in the ascospore wall, providing a convenient means of recognising spore-forming colonies without requiring microscopic examination of every library transformant (Fig. 1A,B). The screening design allowed the mutant and reference populations to be analysed under comparable sporulation conditions, while maintaining the original transformant plate as the source from which individual candidates could subsequently be retrieved. The optical phenotype was therefore used at a stage when colony numbers were too large for exhaustive microscopic scoring, before the sporulation phenotype of any selected candidate was evaluated quantitatively.

**Figure 1.**
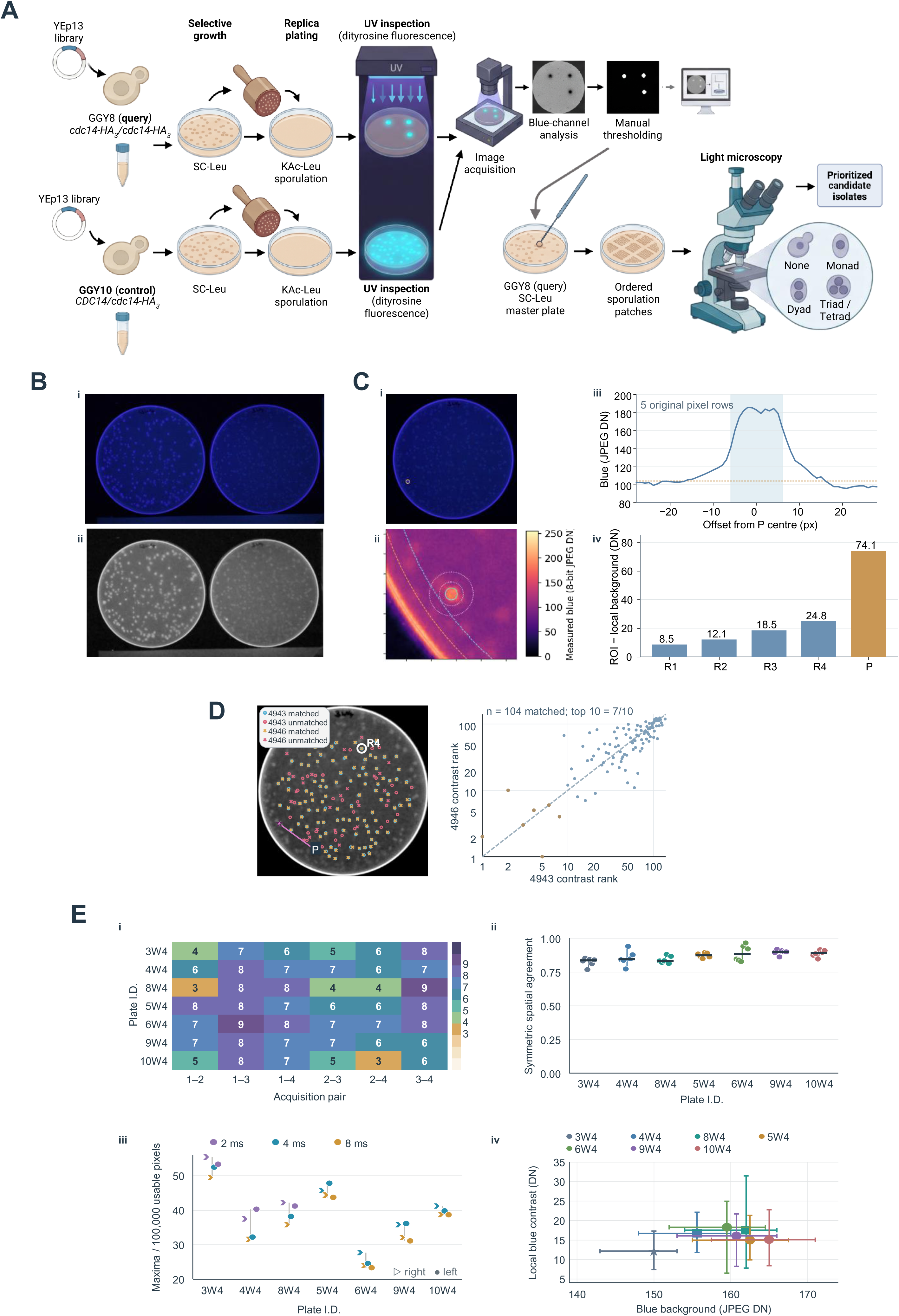
UV-assisted multicopy screening and computational fluorescence analysis. (A) Experimental workflow linking YEp13 genomic-library transformation, sporulation induction, UV fluorescence inspection, colony recovery and microscopy. Panel A was created in BioRender. Carballo, J. (2026) https://BioRender.com/7gvm3pp. (B) Representative original paired-plate UV acquisition showing homozygous *cdc14-HA3* query and sporulation-proficient heterozygous reference material. (C) Blue-channel fluorescence imaging, pseudocolour representation and local intensity measurements illustrating the characterisation of a peripheral fluorescence-associated feature. (D) Spatial detection and correspondence of fluorescence-associated maxima between repeated views. (E.i–iv) Cross-plate quantitative assessment of computational UV-fluorescence detection: strict top-ten overlap, symmetric spatial agreement, detection density per eligible area and comparisons of local signal/background measures. Candidate detections represent image features rather than microscopically verified colony identities; additional image examples and quality-control analyses are shown in Supplementary Fig. S1 and S2.

UV photographs revealed fluorescent colonies against the background of the sporulation plates. The query and sporulation-proficient reference plates were photographed together under controlled UV illumination, using bracketed exposures to retain localised fluorescent signals while limiting background intensity. Additional acquisitions were obtained after exchanging the positions of the paired plates, allowing fluorescence-associated regions to be assessed under different positional conditions without relying on their appearance in a single photograph. A representative paired acquisition shows the mutant query and the sporulation-proficient control (Fig. 1B). The fluorescent areas appeared as discrete, locally bright regions within the photographed colony fields. Inspection of the original paired-plate images was retained as an essential component of the selection procedure, because the physical position of each candidate had to be related to the corresponding colony on the master plate. Photographing both the mutant and sporulation-proficient reference also maintained a visual comparison within each acquired field, even when overall exposure and background levels were not identical between plate arrangements.

Separation of the photographic colour channels showed that the blue channel provided the clearest contrast between localised fluorescence and the surrounding background. Image-specific threshold adjustment facilitated the visual identification of fluorescent regions, which were checked against the original colour photographs and the corresponding source plates (Fig. 1A). The blue-channel signal and local intensity measurements are illustrated separately in Fig. 1C. This procedure supported the selection of UV-positive colonies without imposing a universal fluorescence cutoff or using image intensity as a direct quantitative measure of sporulation.

The selected positions were identified on the corresponding SC-Leu source plates, which had been retained for colony recovery, and the recovered colonies were regrown and patched onto fresh sporulation medium. Microscopy was used to measure sporulation in the recovered colonies. Across six rounds of transformation, approximately 14,400 colonies were inspected by UV illumination. The recovered isolates were retained for quantitative phenotyping and subsequent genetic analysis.

### Computational image analysis extends fluorescence detection across screening plates

To assess the reproducibility of fluorescence detection in the archived UV photographs, we developed a computational procedure that identifies local blue-channel intensity maxima and compares their positions and relative intensities between repeated images of the same plate (Fig. 1D). Each detection was evaluated against its surrounding background within an eligible plate region, and corresponding features were identified after spatial registration. This allowed two related but distinct properties to be examined: whether fluorescence-associated features were found repeatedly at the same positions, and whether those features maintained similar intensity rankings across acquisitions. Unlike the image processing used for the original visual selection of candidates (Fig. 1C), this retrospective analysis provided quantitative measures of detection repeatability, without assigning detected maxima to historical colony identities. The assessment of a peripheral fluorescence signal is illustrated in Supplementary Fig. S1, while Supplementary Fig. S2A provides an example of how detections were matched between photographs.

The procedure was developed using four photographs of plate 3W4 (see Methods for plate nomenclature) and then tested, without retuning the detector, on six additional GGY8 plate families. Four contained previously identified candidate colonies (5W4, 6W4, 9W4 and 10W4), while 4W4 and 8W4 were included as further comparison plates, independently of their observed fluorescence. These comparison plates were not treated as fluorescence-negative controls. Each of the six additional plates was represented by four original photographs, yielding 24 acquisitions and 36 within-plate image pairs; the four development photographs supplied another six pairs. Exposure and plate position differed between acquisitions. The six comparisons within each plate were retained for analysis, but repeated photographs were technical observations, not independent biological replicates. The validation therefore spanned different photographic conditions without selecting plates because their UV signals appeared especially favourable. Thus, consistency was assessed plate by plate rather than by treating every photographic comparison as a separate screen. Representative image-registration, plate-geometry and detection quality-control examples from the 3W4 and 4W4 plates are shown in Supplementary Fig. S2B,C.

We first compared the ten highest-ranked fluorescence-associated detections between pairs of photographs (Fig. 1E.i). The median strict top-ten overlap ranged from 5.5 to 7.5 across the six additional plate families. In the order 4W4, 8W4, 5W4, 6W4, 9W4 and 10W4, the respective medians were 7, 6, 7.5, 7.5, 7 and 5.5 shared detections. Individual image pairs shared between three and nine of their ten highest-ranked maxima; for 10W4, one comparison retained only three. These results indicate that the relative prominence of local signals could change appreciably between photographs. A feature need not retain a top-ten rank, however, to remain detectable at the corresponding plate position. This was particularly relevant for 10W4, whose strongest-signal rankings showed the lowest median agreement in the set.

We therefore examined spatial correspondence independently of intensity ranking (Fig. 1E.ii). Following image registration, median symmetric agreement between detections ranged from 83.2% to 89.9% across the six additional families. Among candidate-containing plates, median agreement was approximately 87% for 5W4, 88% for 6W4, 90% for 9W4 and 89% for 10W4. The additional comparison plates showed approximately 84% agreement for 4W4 and 83% for 8W4. All 42 within-plate registrations, including the six pairs from the 3W4 development plate, met the predefined geometric diagnostics. The agreement remained high across plates with different numbers of detected features, indicating that most localised positions could be recovered in repeated images even when their brightness ranks changed. Passing the geometric diagnostics supported comparison of corresponding regions but did not establish the biological identity of any signal.

Detection abundance varied among plates and acquisitions (Fig. 1E.iii). Across four photographs per plate, 5W4 yielded 103–113 eligible maxima, 6W4 yielded 55–66, 9W4 yielded 73–85 and 10W4 yielded 91–98; the corresponding ranges were 75–95 for 4W4 and 85–100 for 8W4. After normalisation to 100,000 eligible image pixels, median detection density ranged from 24.3 for 6W4 to 45.0 for 5W4 among the candidate-containing plates, while 4W4 and 8W4 yielded medians of 34.8 and 39.7, respectively. They must not be interpreted as percentages of sporulating colonies: individual maxima were not linked to microscopically scored colonies, and the total number of colonies within each analysed image was not established.

Local signal and surrounding-background measurements provided an additional assessment of fluorescence-associated maxima across acquisitions obtained under different photographic conditions (Fig. 1E.iv). These measurements complemented the positional and intensity-ranking analyses by characterising the local contrast of detected features. Overall, spatial correspondence was more consistent than the ranking of the strongest signals, although detection abundance and local contrast varied between photographs. The computational analysis thus established reproducibility at the level of image features, complementing the fluorescence-guided selection of colonies performed in the original screen.

### Microscopic quantification identifies a spectrum of improved sporulation phenotypes

The recovered isolates were therefore grown as patches on fresh sporulation medium and assessed directly by light microscopy. The observations distinguished unsporulated units from monads, dyads, triads and tetrads, allowing the sporulation phenotype to be described both as total ascus formation and as the production of asci containing two or more spores. Total sporulation included monads, dyads, triads and tetrads, whereas partial sporulation counted dyads, triads and tetrads but excluded monads. Monads contributed variably to the total sporulation values. Partial sporulation was therefore used to rank recovered candidates. In particular, a colony with numerous monads could show a relatively high total-sporulation value while having substantially lower partial sporulation. Retaining both readouts allowed the selected colonies to be characterised across different stages of spore formation, rather than reducing the microscopic observations to a single fluorescence-associated score.

The corrected plate-assignment key identified 739 microscopy records from colonies selected during the six transformation rounds, comprising 617 GGY8 query records and 122 GGY10 reference records. These observations represent the subset recovered for microscopic assessment from the approximately 14,400 colonies initially inspected by UV fluorescence. Figure 2A displays the distribution of total sporulation across both the mutant and control records, illustrating the broad range of ascus formation observed among the selected colonies.

**Figure 2.**
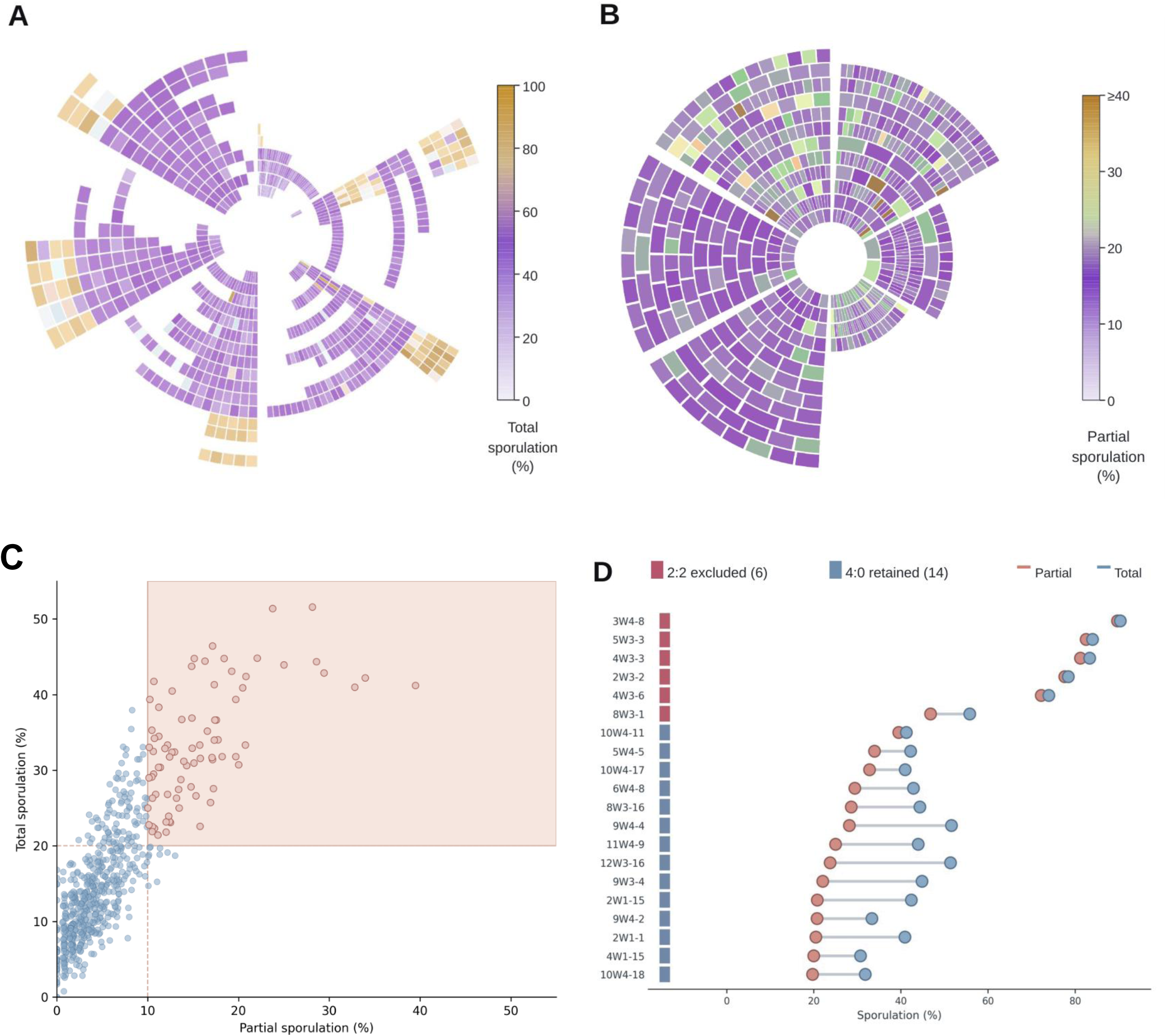
Quantification and genetic disposition of candidates from the primary screen. (A) Total sporulation among 739 selected-colony records, comprising 617 homozygous mutant query and 122 heterozygous control records. (B) Partial sporulation in the 617 corrected query records; the displayed colour scale is limited at 40% for visual differentiation without changing higher underlying observations. (C) Scatter relationship between partial sporulation on the X axis and total sporulation on the Y axis. The shaded region is illustrative, not an experimentally documented universal cutoff. (D) Total and partial sporulation values for the top-20 corrected query colonies, indicating six isolates with 2:2 G418 segregation (red) and 14 with the 4:0 pattern consistent with the intended mutant-marker background (blue). These records represent the selected subset subjected to microscopy, not all approximately 14,400 colonies surveyed by UV.

Partial sporulation was then examined specifically within the 617 corrected GGY8 query records (Fig. 2B). This representation excludes monads from the sporulation numerator and therefore distinguishes colonies producing asci with two or more spores from those whose total sporulation was elevated predominantly by monad formation. The colour scale is limited at 40% for visual discrimination, without changing the underlying measurements.

Plotting partial sporulation against total sporulation for the mutant records further illustrates the relationship between the two measurements (Fig. 2C). Colonies with similar total-sporulation values could differ substantially in partial sporulation, consistent with variation in the contribution of monads. The shaded region is illustrative and does not represent a retrospectively established universal selection threshold.

The 20 query colonies with the highest corrected partial-sporulation values were carried forward as the primary-hit set. Subsequent genetic and plasmid-based assessment distinguished retained candidates from isolates whose increased sporulation did not provide suitable evidence of suppression in the intended mutant background (Fig. 2D).

### Independent plasmid retransformation identifies reproducible suppressors

To determine whether their unusually high sporulation was plasmid-associated, the six most strongly sporulating primary isolates (3W4-8, 5W3-3, 4W3-3, 2W3-2, 4W3-6 and 8W3-1) were first tested by retransformation of the recovered plasmids into fresh GGY8 recipients. The six plasmids failed to reproduce the elevated sporulation observed in their original isolates. In contrast, the original isolates retained high sporulation when recovered from frozen stocks and scored again (Supplementary Fig. S3). Subsequent tetrad analysis revealed 2:2 G418-resistance segregation in all six isolates, rather than the 4:0 pattern expected for the homozygous *cdc14-HA3* background. These six isolates were consequently excluded from the retained primary-hit set, leaving 14 isolates for further assessment (Fig. 2D).

Five plasmids from the 14 genetically retained primary isolates—10W4-11, 9W4-4, 10W4-17, 5W4-5 and 6W4-8—were selected for quantitative assessment following independent introduction into fresh GGY8 recipients (Fig. 3A). Four independent transformants were examined for each candidate plasmid, together with eight empty-vector transformants. Sporulation was scored in repeated assays, and each independent transformant contributed one summary value to the statistical comparisons. This design allowed plasmid-associated effects to be evaluated against contemporaneous empty-vector controls rather than relying on the unusually high sporulation of the initially selected colonies.

**Figure 3.**
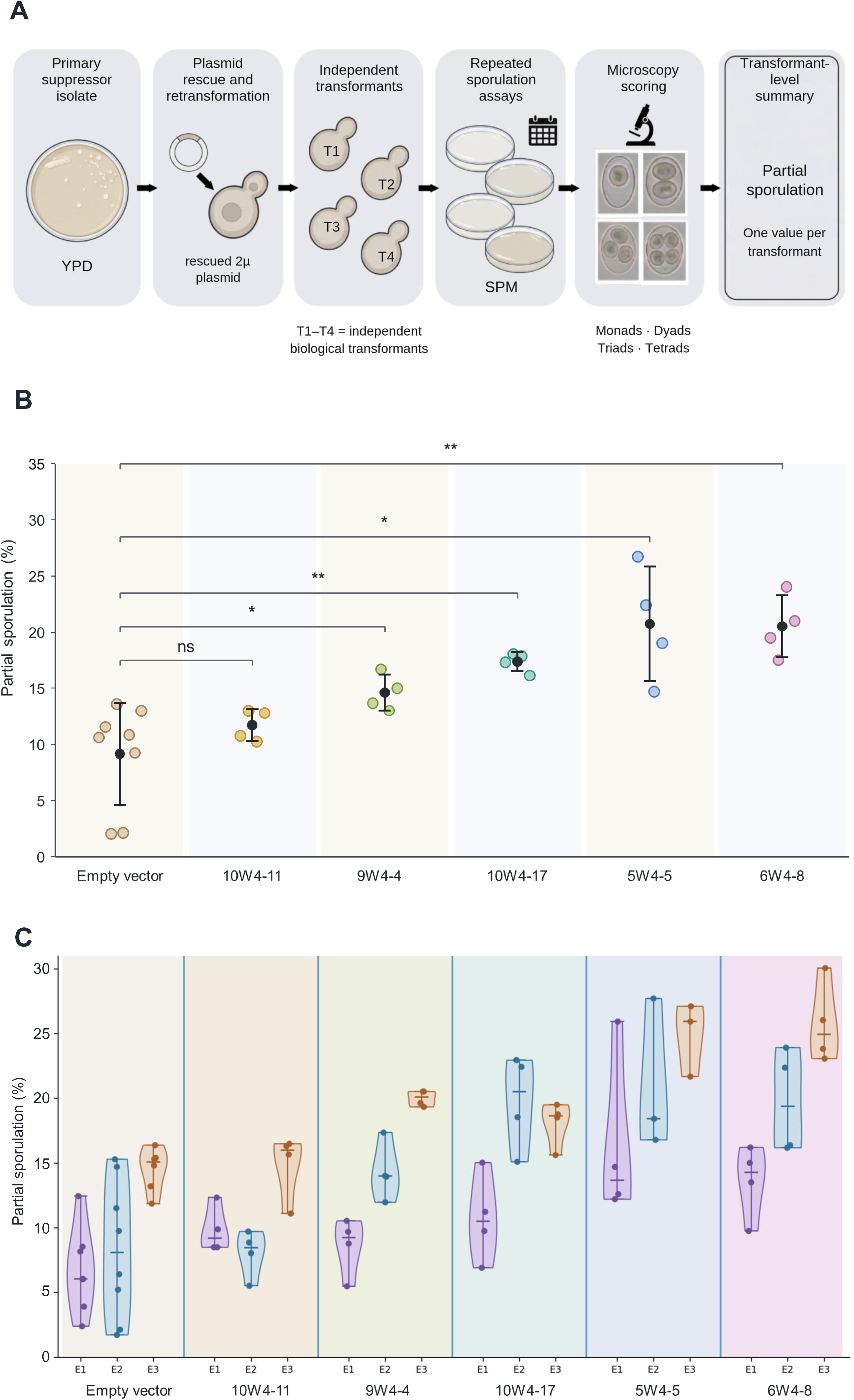
Reproducibility of multicopy suppression following independent plasmid retransformation. (A) Experimental design for independent plasmid introduction and repeated sporulation assays. This panel was created in BioRender. Carballo, J. (2026) https://BioRender.com/8k3sh7o. (B) Transformant-level partial-sporulation summaries for the five candidate plasmids and empty-vector control; each independent transformant contributes one inferential observation. (C) Experiment-resolved E1–E3 violin plots showing repeated assay observations as individual points. The violin shapes provide a smoothed view of the spread of these measurements. Statistical comparisons were based on independent transformants, not repeated observations. For each candidate n = 4 and for empty-vector control n = 8. Five candidate-versus-vector Welch tests and exact studentised-permutation comparisons were each Holm-adjusted across the five comparisons; values appear in Table 1. Brackets indicate comparisons with empty vector; significance symbols correspond to Holm-adjusted Welch-test P values (ns, P ≥ 0.05; *P < 0.05; **P < 0.01).

**Table 1.**
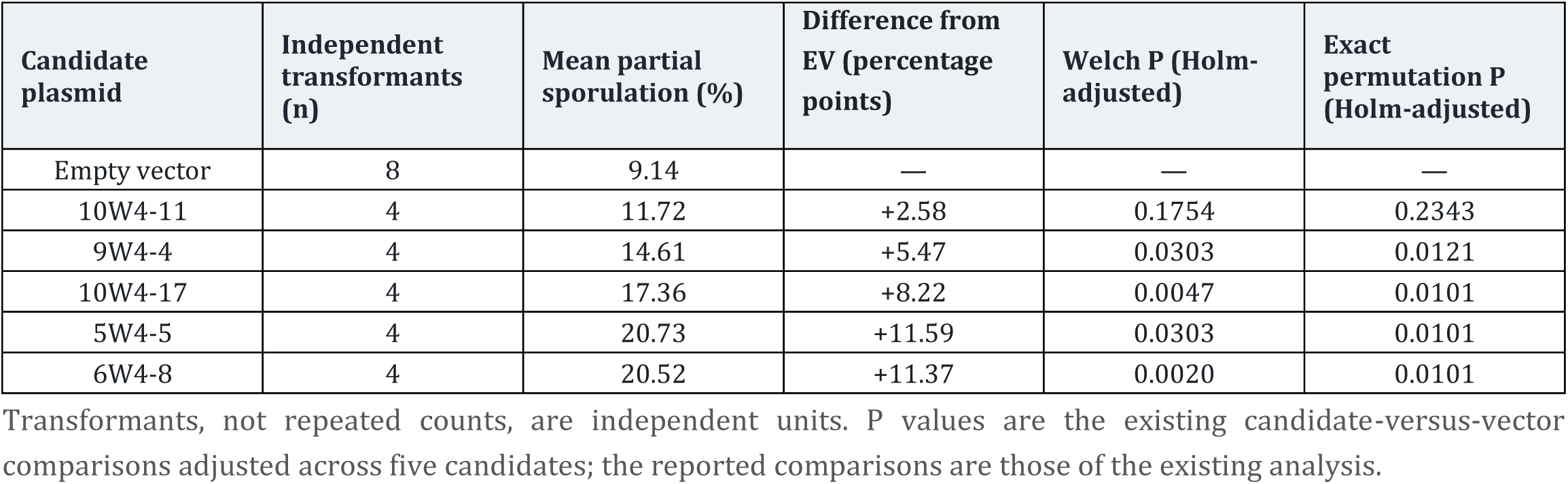
Independent retransformation: transformant-level partial sporulation.

| **Candidate plasmid** | **Independent transformants (n)** | **Mean partial sporulation (%)** | **Difference from EV (percentage points)** | **Welch P (Holm-adjusted)** | **Exact permutation P (Holm-adjusted)** |
|---|---|---|---|---|---|
| Empty vector | 8 | 9.14 | — | — | — |
| 10W4-11 | 4 | 11.72 | +2.58 | 0.1754 | 0.2343 |
| 9W4-4 | 4 | 14.61 | +5.47 | 0.0303 | 0.0121 |
| 10W4-17 | 4 | 17.36 | +8.22 | 0.0047 | 0.0101 |
| 5W4-5 | 4 | 20.73 | +11.59 | 0.0303 | 0.0101 |
| 6W4-8 | 4 | 20.52 | +11.37 | 0.0020 | 0.0101 |

Transformants, not repeated counts, are independent units. P values are the existing candidate-versus-vector comparisons adjusted across five candidates; the reported comparisons are those of the existing analysis.

All five plasmids produced higher mean partial-sporulation efficiencies than the empty-vector reference of 9.1%, although only four increases were supported by the adjusted statistical comparisons (Fig. 3B; Table 1). The highest means were obtained with 5W4-5 (20.7%) and 6W4-8 (20.5%), followed by 10W4-17 (17.4%) and 9W4-4 (14.6%). In contrast, 10W4-11, despite its strong primary-screen phenotype, yielded a mean of 11.7% following independent retransformation. The four candidate-versus-vector comparisons involving 5W4-5, 6W4-8, 10W4-17 and 9W4-4 were supported by the existing Holm-adjusted Welch tests and exact studentised-permutation comparisons. The increase associated with 10W4-11 was not supported after adjustment (Table 1).

The repeated assays also revealed variation between experimental series E1–E3 (Fig. 3C). These experiment-resolved distributions show how the sporulation measurements varied within and between the tested transformants, including the greater overlap between 10W4-11 and empty-vector observations. The experimental series provided repeated observations of the same independently established transformants and were not counted as additional biological replicates in the inferential analysis.

The partial-sporulation efficiencies obtained after retransformation were generally lower than those recorded for the corresponding original primary isolates. The two stages nevertheless addressed different questions: the initial screen identified unusually sporulating colonies for further examination, whereas the independent retransformation assays tested whether their recovered plasmids produced a consistent increase relative to empty-vector controls across fresh recipients and repeated measurements. The available data do not establish a single cause for the differences in magnitude between the primary and follow-up measurements. Taken together, the independent assays support plasmid-associated suppression for four of the five candidates examined, while 10W4-11 remains an example of a strong initial phenotype that was not confirmed by the quantitative follow-up.

### Genomic architecture reveals biologically distinct suppressor-associated regions

The three plasmids showing the largest mean increases in partial sporulation following independent retransformation—5W4-5, 6W4-8 and 10W4-17—were characterised using their archived end-sequencing and genomic mapping records (Fig. 4A). The inserts mapped to different chromosomal intervals and contained distinct combinations of complete, truncated and dubious annotated ORFs.

**Figure 4.**
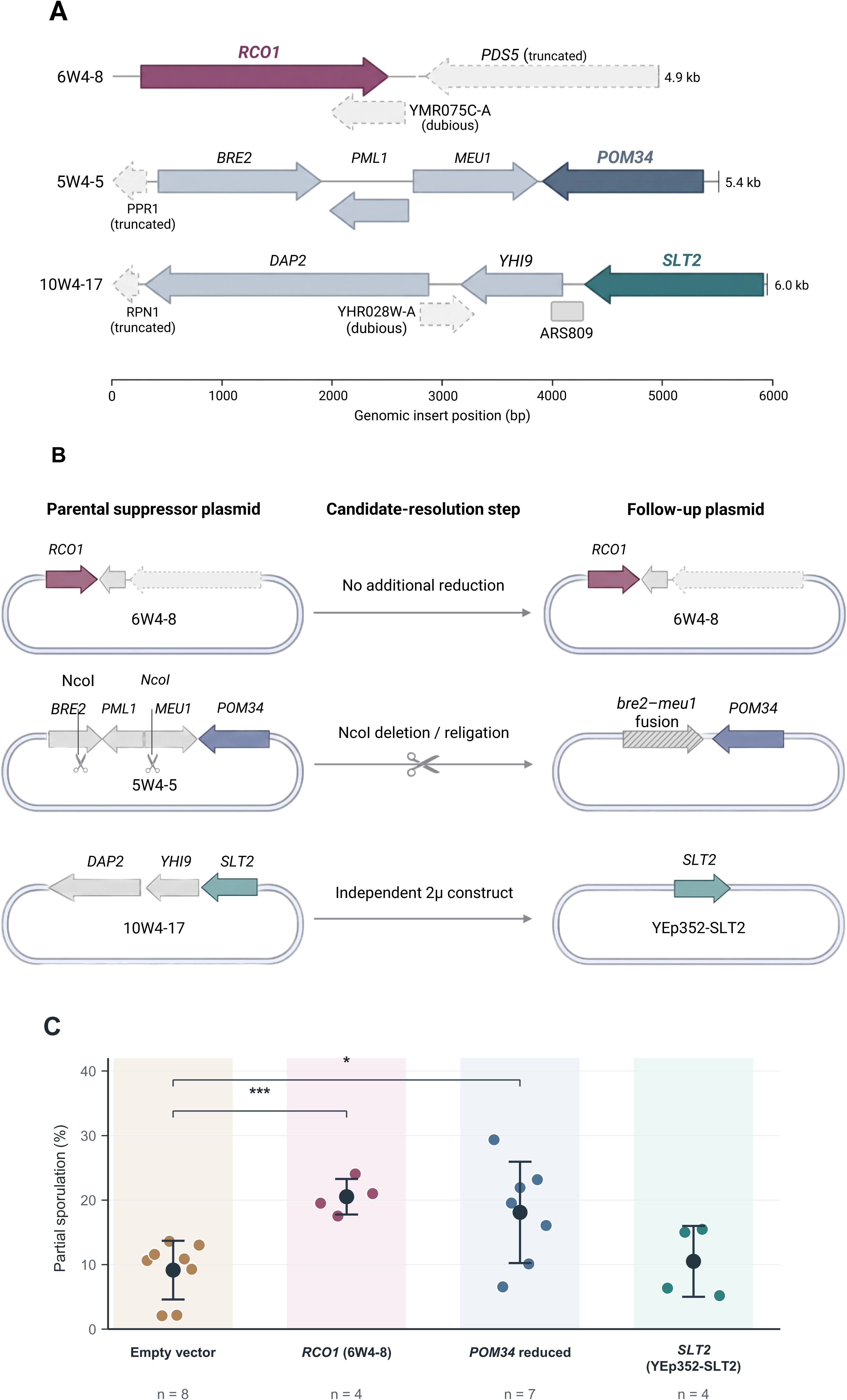
Genomic organisation of suppressor-associated fragments and molecular follow-up of single-ORF candidate plasmids. (A) Mapped fragments 6W4-8 (4,916 bp), 5W4-5 (5,433 bp) and 10W4-17 (6,024 bp), including annotated intact, dubious and truncated features. (B) Candidate interpretation based on the original 6W4-8/*RCO1* fragment, the *POM34* single-ORF plasmid obtained through NcoI deletion/religation of the 5W4-5 insert and the independently acquired *SLT2* single-ORF plasmid. The *POM34* molecular product also retains rearranged *bre2*–*meu1* sequence. (C) Partial sporulation of the empty-vector reference (n = 8), *RCO1* (n = 4), *POM34* (n = 7) and *SLT2* (n = 4), with n indicating independent transformants. The 6W4-8 data are the same transformant-level observations presented in Fig. 3. Table 2 provides the existing candidate-focused comparisons. Brackets indicate comparisons with empty vector; significance symbols correspond to unadjusted Welch-test P values (*P < 0.05; ***P < 0.001). No formal statistical comparison was performed for *SLT2*.

**Table 2.**
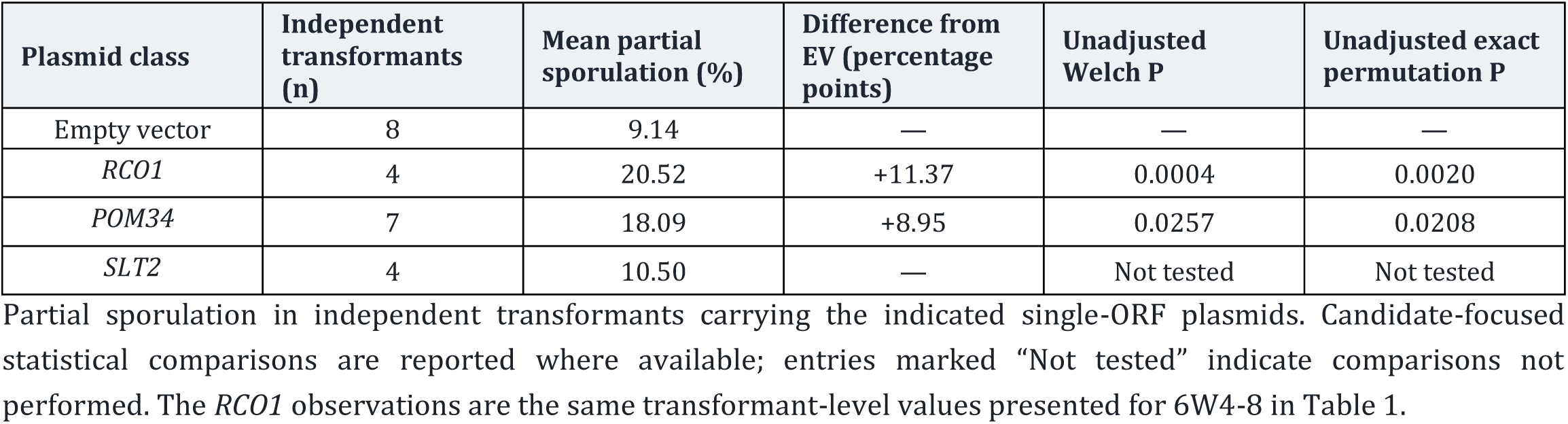
Sporulation of transformants carrying single-ORF plasmids.

| Plasmid class | Independent transformants (n) | Mean partial sporulation (%) | Difference from EV (percentage points) | Unadjusted Welch P | Unadjusted exact permutation P |
|---|---|---|---|---|---|
| Empty vector | 8 | 9.14 | — | — | — |
| *RCO1* | 4 | 20.52 | +11.37 | 0.0004 | 0.0020 |
| *POM34* | 7 | 18.09 | +8.95 | 0.0257 | 0.0208 |
| *SLT2* | 4 | 10.50 | — | Not tested | Not tested |

The 6W4-8 plasmid carried a 4,916-bp chromosome XIII fragment containing intact *RCO1*, a truncated portion of *PDS5* and the dubious annotated feature *YMR075C-A*. *RCO1* was the only intact conventional candidate ORF within this interval.

The 5W4-5 plasmid contained a 5,433-bp fragment from chromosome XII, encompassing the complete *POM34*, *PML1*, *BRE2* and *MEU1* coding sequences together with a truncated *PPR1* region. Thus, unlike 6W4-8, this recovered fragment contained several intact candidate ORFs.

The 10W4-17 plasmid contained a 6,024-bp chromosome VIII fragment with intact *DAP2*, *YHI9* and *SLT2* ORFs, a truncated *RPN1* region, the dubious feature *YHR028W-A* and the annotated *ARS809* element. The presence of multiple intact ORFs in 5W4-5 and 10W4-17 prevented assignment of their suppression phenotypes to individual genes on the basis of the parental plasmids alone.

### Molecularly defined plasmids retain or distinguish suppressor activity

To investigate which genes might contribute to suppression, we focused on the three genomic-library plasmids with the largest reproducible increases in partial sporulation. Candidate ORFs were prioritised on the basis of their annotated cellular functions and potential connections with meiotic regulation. We selected *RCO1* from 6W4-8, *POM34* from 5W4-5 and *SLT2* from 10W4-17 for a more restricted molecular assessment (Fig. 4B). *RCO1* encodes a component of the Rpd3S histone-deacetylase complex involved in chromatin-associated transcriptional regulation (Carrozza et al., 2005). *POM34* encodes an integral membrane component of the nuclear pore complex (Miao et al., 2006), whereas *SLT2* encodes a MAP kinase involved in cell-wall-integrity signalling (Martín et al., 1993). These functional associations provided reasons to prioritise the three candidates, without establishing that any was a direct substrate of Cdc14.

The original 6W4-8 fragment already contained *RCO1* as its only intact conventional candidate ORF and was therefore retained without further reduction (Fig. 4B). Its mean partial-sporulation efficiency was 20.5%, compared with 9.1% for the empty-vector reference (Fig. 4C; Table 2). These are the same independent-transformant observations reported for 6W4-8 in Fig. 3B and Table 1, rather than an additional set of experiments.

For *POM34*, a reduced plasmid was generated from the multigenic 5W4-5 fragment by NcoI digestion and religation. The resulting product retained *POM34* as its only intact conventional candidate ORF, although the rearranged insert also contained a *bre2–meu1* fusion (Fig. 4B). Seven independent transformants carrying this *POM34* single-ORF plasmid gave a mean partial-sporulation efficiency of 18.1%, compared with 9.1% for the empty-vector reference (Fig. 4C; Table 2). The existing unadjusted Welch (P = 0.026) and exact studentised-permutation (P = 0.021) comparisons supported the increase. Its mean was also close to that observed for the parental 5W4-5 plasmid (20.7%), although the parental and reduced constructs were tested in separate retransformation series rather than side by side. Suppression was therefore retained after removal of several intact ORFs from the original fragment.

Finally, an independently obtained high-copy *SLT2* single-ORF plasmid was tested as a candidate corresponding to the 10W4-17 insert (Fig. 4B). The *SLT2* construct yielded a mean partial-sporulation efficiency of 10.5% across four independent transformants (Fig. 4C; Table 2), whereas the original multigenic 10W4-17 plasmid yielded 17.4% in the separate retransformation series (Fig. 3B; Table 1). Unlike the *RCO1*-associated fragment and reduced *POM34* plasmid, the independently tested *SLT2* construct did not reproduce the enhanced sporulation of the genomic-library plasmid. These results do not identify which component of 10W4-17 accounts for suppression, but indicate that *SLT2* alone, in the construct and conditions examined, was insufficient to recapitulate the original plasmid-associated response.

## Discussion

The suppression of the *cdc14-HA3* sporulation defect by increased gene dosage provides a genetic approach to processes that contribute to successful meiotic progression. In our earlier analysis, reduced Cdc14 activity affected the processing of meiotic recombination intermediates, Yen1 regulation and subsequent chromosome segregation (Alonso-Ramos et al., 2021). The sporulation phenotype of the mutant is therefore interpreted in the context of the recombination and chromosome-segregation defects characterised previously. Multicopy suppression allowed us to search beyond the established Cdc14 and FEAR/MEN regulators and identify genomic regions that enhanced the final meiotic phenotype.

UV-excited dityrosine fluorescence enabled an initial survey of approximately 14,400 library transformants for colonies producing spores, with microscopy and genetic analysis subsequently used to identify candidates for plasmid testing. The combination of original photographic acquisitions with image processing was particularly useful for retaining the positions of candidate colonies on recoverable source plates. The screening procedure therefore makes it possible to examine a complex meiotic phenotype at library scale without relying on a selectable colony-growth advantage (Briza et al., 1986). Such an assay is particularly useful for meiotic mutants whose defects are manifested after entry into the sporulation programme and cannot be selected conveniently by increased colony proliferation. The two-stage approach allowed sporulation-associated fluorescence to identify candidates and subsequent microscopy to compare their ascus phenotypes. Photographic storage of the UV plates also preserved experimental material for analyses beyond the initial manual selection, providing the basis for the computational evaluation performed here.

Computational analysis of the archived UV photographs showed that fluorescence-associated features could be located reproducibly across repeated acquisitions, even when their relative intensity rankings varied. This analysis was performed retrospectively, and the computationally detected maxima were not assigned to individual colonies subsequently validated as suppressors. The results therefore establish the repeatability of image-based fluorescence detection rather than automated identification of biological suppressors. Nevertheless, the procedure offers a promising basis for further automation of similar genomic screens. Linking computational detections to individual colonies and their microscopic sporulation phenotypes could facilitate more extensive candidate recovery and quantitative assessment in future applications.

Microscopic scoring identified considerable variation in ascus production among colonies selected by UV fluorescence. Total sporulation and partial sporulation distinguished the production of monads from asci containing multiple spores, while genetic analysis established which isolates retained the expected mutant marker configuration. This assessment produced 14 retained isolates from the 20 leading primary hits (Fig. 2D). The subsequent plasmid tests identified genomic regions that reproducibly enhanced sporulation in fresh *cdc14-HA3* recipients.

Four of the five tested plasmids increased partial sporulation in independently transformed recipients, with the two strongest, 5W4-5 and 6W4-8, producing means more than twice that of the empty-vector reference. These are substantial changes for a mutant in which efficient spore formation is severely compromised. Moreover, the supported responses were distributed across plasmids with different genomic content, broadening the set of molecular contexts potentially associated with improved meiosis. The Welch and exact studentised-permutation comparisons supported the four positive plasmid responses at the transformant level. The additional supported responses of 10W4-17 and 9W4-4 extended the range of genomic regions associated with improved ascus formation. This degree of reproducibility is especially informative because the original selection was based on the sporulation of individual colonies, whereas the subsequent evaluation used fresh transformants carrying independently recovered plasmids.

The molecular composition of the recovered suppressor-associated fragments points to two particularly interesting biological contexts: chromatin regulation and nuclear organisation. The 6W4-8 insert provides a particularly informative link to chromatin regulation because *RCO1* is its sole intact conventional candidate ORF. Rco1 is a subunit of the Rpd3S histone-deacetylase complex. Its PHD1 domain preferentially recognises unmodified H3K4, whereas the Eaf3 chromodomain recognises H3K36-methylated nucleosomes. Together, these interactions contribute to Rpd3S targeting and the suppression of inappropriate transcription initiation within coding regions (Carrozza et al., 2005; Keogh et al., 2005; Li et al., 2007). Rpd3 complexes also regulate the meiotic transcriptional programme: Rpd3L and Rpd3S have distinct effects on the expression of meiosis-specific regulators, including *IME1*, *IME2* and *NDT80* (Yeheskely-Hayon et al., 2013). These established functions make a dosage-sensitive connection between chromatin-associated transcription and the *cdc14-HA3* meiotic phenotype biologically plausible. The strong suppression conferred by 6W4-8 thus prioritises the *RCO1*-associated region for further investigation without requiring a direct biochemical interaction between Rco1 and Cdc14. The 6W4-8 result makes it possible to consider this chromatin-associated activity in a sensitised *CDC14* background, while the mechanism underlying suppression remains an open experimental question.

The *POM34*-containing region introduces nuclear-pore organization as another potentially relevant biological context for suppression. Pom34 is an integral membrane nucleoporin associated with nuclear pore complex assembly (Miao et al., 2006; Onischenko et al., 2009). Nuclear pore complexes undergo extensive remodelling during budding-yeast meiosis, including the redistribution of core and nuclear-basket components (King et al., 2019; King et al., 2023). Pom34 itself has been followed during the meiotic sequestration and selective inheritance of nuclear pore complexes (Koch et al., 2020; Ruediger et al., 2026). The persistence of suppressor activity in the reduced *POM34* single-ORF plasmid makes this region particularly attractive for further analysis. An additional consideration is the potential phosphoregulation of Pom34: several phosphorylation sites have been identified in vivo, including a site associated with Cdk1-dependent phosphorylation (Holt et al., 2009). Whether these modifications contribute to the meiotic functions of Cdc14 or to the observed suppression remains to be established. The reduced plasmid also contains a *bre2–meu1* chimeric sequence, absent as such from the parental construct, whose possible contribution cannot yet be excluded. We are currently pursuing further molecular analysis of the *POM34*-associated region to distinguish these possibilities.

Together, these findings demonstrate that a UV-assisted multicopy screen can recover genomic fragments capable of reproducibly improving sporulation in a sensitised meiotic mutant. Independent retransformation and molecular characterisation extended the original screen beyond fluorescence-based candidate selection, identifying suppressor-associated regions linked to chromatin regulation and nuclear-pore organisation. The accompanying computational analysis provides a quantitative framework for improving fluorescence detection in future screens. By combining genetic screening with independent phenotypic validation and molecular analysis of recovered plasmids, this work establishes a useful experimental foundation for investigating how increased gene dosage can compensate for defects in Cdc14-dependent meiotic progression.

## Materials and Methods

### Yeast strains, growth conditions and multicopy library

The screen used SK1-derived diploid GGY8, homozygous for the C-terminally tagged *cdc14-HA3* allele, and the sporulation-proficient heterozygous control GGY10 (*CDC14*/*cdc14-HA3*). The genotypes of all strains are listed in Supplementary Table S1A. The tagged allele is referred to as *cdc14-HA* in Alonso-Ramos et al. (2021).

The genomic library was based on the high-copy YEp13 vector carrying LEU2. Partially digested AB320 yeast genomic DNA was cloned at the YEp13 BamHI site (Broach et al., 1979; Nasmyth and Tatchell, 1980). Plasmids and mapped genomic-library inserts are listed in Supplementary Table S1B. The screen yielded 20 primary candidates, of which 14 were retained for further analysis following genetic assessment. Five plasmids (10W4-11, 9W4-4, 10W4-17, 5W4-5 and 6W4-8) were included in the independent quantitative retransformation assays. The 20 primary candidates and their recorded follow-up are listed in Supplementary Table S2. The multicopy *cdc14-HA3* plasmid pPD.2, originally described by Jaspersen et al. (1998), was used as a reference for dosage responsiveness, as in Alonso-Ramos et al. (2021).

### Library transformation and induction of sporulation

Yeast transformation followed a lithium-acetate, single-stranded carrier-DNA and polyethylene-glycol procedure (Gietz and Schiestl, 2007). The library transformation mixture contained 2,400 µl of 50% PEG 3500, 360 µl of 1 M lithium acetate, 500 µl carrier single-stranded DNA (2 mg/ml), 100 µl of the YEp13 library and 240 µl water. Cells were plated onto SC-Leu and grown at 30°C for approximately 72 h. Colonies were replica-plated onto leucine-deficient potassium-acetate sporulation medium (KAc-Leu) and incubated for approximately 72–96 h at 30°C. The original SC-Leu plates were maintained at approximately 4°C for subsequent colony retrieval. Six library-transformation rounds were performed.

Solid KAc-Leu medium contained 2% (w/v) potassium acetate, 0.22% yeast extract, 0.05% (w/v) glucose, 0.087% amino-acid mixture excluding leucine and 2.5% agar.

### UV fluorescence and photographic acquisition

Sporulation plates were placed face-down, with the agar surface uppermost, on a High Performance UV Transilluminator (UVP) fitted with 25-W UV tubes. The UV illumination was operated at its lowest power setting. Two plates, one containing the mutant query and the other the sporulation-proficient control, were positioned with a card guide within a region of comparatively uniform illumination. A camera mounted at fixed height on an overhead support photographed both plates. For paired acquisitions, the positions of the query and reference plates were exchanged before obtaining an additional bracketed series.

A Canon EOS 1100D with a 22.2 × 14.7 mm CMOS sensor was used. Acquisition was performed in manual (M) mode, 1/250-s shutter speed, f/5.6 aperture, ISO 6400, auto-exposure bracketing (AEB) of −1, 0 and +1, neutral picture style, tungsten white balance (∼3200 K), standard auto-lighting optimiser, partial metering, one-shot autofocus and a 2-s self-timer. No flash was used. Three-frame bracketed sequences were acquired for paired plates. Archived images used in the computational analysis had EXIF exposure times of 2, 4 or 8 ms (f/5.6, ISO 6400).

### Image processing and candidate selection

Photographs were cropped to retain both plates and their circular boundaries. The Fiji distribution of ImageJ was used to separate red, green and blue channels and inspect localised UV-excited signals (Schneider et al., 2012; Schindelin et al., 2012). The blue channel provided useful signal-to-background contrast. Intensity thresholds were adjusted for each image, and processed photographs were compared with the corresponding original colour images and source plates. No universal fluorescence threshold was applied across all plates.

Selected UV-positive colonies were recovered from the matching SC-Leu plates, transferred to fresh SC-Leu and regrown for approximately 2 days at 30°C. They were patched onto KAc-Leu in a grid with GGY10 control patches at the corners and incubated for 72–96 h. Microscopic sporulation scoring provided the primary quantitative phenotype, followed where documented by repeat UV examination of the patches. This second examination was an auxiliary check of fluorescence–sporulation correspondence, not a substitute for the microscopic count.

### Computational analysis of UV-fluorescence images

The computational detection and registration procedure was developed on four photographs from the 3W4 reference plate and applied without detector retuning to six further GGY8 plate families (4W4, 5W4, 6W4, 8W4, 9W4 and 10W4), each photographed four times. The 3088 × 2056 RGB JPEGs were resized once to 2048 × 1363 pixels using Lanczos interpolation, without gamma, exposure, flat-field or intensity correction. Candidate fluorescence-associated positions were identified in the blue channel by a difference-of-Gaussians filter (σ = 2 and 18 pixels), a 21-pixel local-maximum window, a score threshold above 4 digital numbers and a minimum 22-pixel spacing. Eligible centres were evaluated within the inner 90% plate ellipse, with local contrast measured from a radius-6-pixel signal region against an annulus of 12–22 pixels. Repeated photographs of the same plate were aligned using central-texture rotation/translation registration, and one-to-one spatial matches were assigned within 8 target-image pixels. Strict top-ten overlap and symmetric spatial agreement were calculated separately. Four photographs and six acquisition pairs for each plate were technical/repeated observations, not biological replicates; detected maxima were not assigned to individual historical colony identifiers. The validation used six additional plate families (24 photographs; 36 within-plate comparisons), while 3W4 remained the development/reference family. Custom Python scripts implemented the image-analysis procedure. Input-image hashes, detection outputs, spatial matches and registration diagnostics are preserved in the reproducibility archive.

### Code availability

The custom scripts used for digital UV-fluorescence image analysis may be requested from the corresponding author. A public repository link will be provided when available.

### Microscopic sporulation counts and candidate ranking

Material from each recovered patch was mounted on a slide and examined by light microscopy. At least 100 cell/ascus units per sample were scored and classified each as unsporulated, monad, dyad, triad or tetrad. Total sporulation (%) was calculated as 100 × (monads + dyads + triads + tetrads)/all units scored. Partial sporulation (%) was calculated as 100 × (dyads + triads + tetrads)/all units scored. Monads remained in the denominator of both calculations but contributed only to total sporulation. Individual spores within an ascus were not independent counted units.

Original colony identifiers were retained in the format *x*W*y*-*z*, in which *x* denotes the plate number, W the 1:10 dilution used for screening (50 µl of transformation mixture plus 450 µl of water), *y* the transformation round and *z* the selected colony number. The corrected plate-assignment key treated plates 1–12 of each transformation round as query GGY8 and plates 13–16 as GGY10 controls, except for plates 5–6 in round 1 and 5–8 in round 2, which were GGY10 controls. This key assigned 617 quantified records to the query and 122 to controls. Primary hits were ranked by corrected query partial sporulation; the 20 primary hits were the 20 highest-ranked corrected GGY8 query records by partial sporulation.

### Genetic screening of retained isolates and plasmid mapping

Selected isolates were subjected to tetrad dissection and G418-resistance segregation analysis. A 4:0 pattern was consistent with retention of the intended homozygous tagged mutant-marker configuration, whereas a 2:2 pattern excluded the isolate from the set of suppressors of that specified background. The segregation result was not used to infer a particular molecular rearrangement or revertant mechanism.

Recovered plasmids were propagated in *Escherichia coli* and analysed by Sanger sequencing from vector-derived primers flanking the genomic insert. Primer pBR2 was 5′-GGAGCCACTATCGACTAC-3′ and pBR3 was 5′-TCCCCATCGGTGATGTC-3′. The resulting terminal reads were mapped to the *S. cerevisiae* reference genome, and the intervening genomic region inferred for ORF annotation (Engel et al., 2014). Insert maps were reconstructed from vector-end sequence reads and the corresponding genomic intervals; MboI/Sau3AI-compatible ends were represented in the archived maps. The reduced 5W4-5 product was generated by NcoI deletion and religation and assessed by mapping and an expected-size gel check and sequencing. Product-class labels A and B represent separate preparations, not independently designed molecular constructs. The mapped *POM34* single-ORF product also retained a predicted *bre2*–*meu1* fusion, as annotated in its preserved xDNA map.

The *SLT2* single-ORF plasmid (YEp352-*SLT2*) was a YEp352-derived 2µ vector, containing a reported approximately 2.2-kb SalI–EcoRI fragment including *SLT2* and its flanking expression sequences, with *URA3* selection. The plasmid was kindly donated by Humberto Martín Brieva.

### Independent retransformation, data reduction and statistical comparisons

Independent retransformation assays included four independent transformants for each of five recovered candidate plasmids and eight empty-vector control transformants. Independent *POM34* single-ORF plasmid transformants were analysed as one molecular product category (n = 7). The *SLT2* single-ORF plasmid was examined in four additional transformants. The experimental archive records multiple dates and media conditions (including SPM, KAc-Leu and KAc-Ura). Transformant identity, rather than each microscopy field, technical count or date, defined the independent biological observation.

For each transformant, valid repeated counts on the same date and medium were first averaged to an experiment-condition summary; the independent-transformant value was then calculated as the mean across its available valid experiment-condition summaries. The five tested candidate plasmids were compared against empty-vector controls using Welch two-sample tests. P values were adjusted by the Holm procedure across the five comparisons. Exact transformant-level permutation of the studentised Welch statistic, followed by Holm adjustment across the same comparison family, was used as a complementary inferential approach. The same 6W4-8 data appear in the *RCO1* row of Fig. 4/Table 2, with the original unadjusted candidate-focused P values. The *POM34* single-ORF plasmid was compared with empty vector using the existing unadjusted Welch and exact permutation tests (6,435 exact allocations). The *SLT2* single-ORF plasmid was analysed descriptively; no formal vector-controlled inference was calculated for that construct.

## Acknowledgements

We gratefully acknowledge the late Dr Marco Geymonat for his early intellectual contribution to the conception of this multicopy suppressor screen and for generously sharing genetic material that helped initiate the project. We also thank Dr Humberto Martín Brieva for providing the YEp352-*SLT2* plasmid. We are grateful to colleagues at the Genome Damage and Stability Centre (University of Sussex), where the original screening experiments were conducted, and at the Institute of Functional Biology and Genomics (IBFG) for helpful discussions, reagents and technical assistance.

## References

Acosta I, Ontoso D, San-Segundo PA (2011). The budding yeast polo-like kinase Cdc5 regulates the Ndt80 branch of the meiotic recombination checkpoint pathway. Molecular Biology of the Cell 22, 3478–3490. 10.1091/mbc.E11-06-0482.

Alonso-Ramos P, Carballo JA (2024). Decoding the nucleolar role in meiotic recombination and cell cycle control: insights into Cdc14 function. International Journal of Molecular Sciences 25, 12861. 10.3390/ijms252312861.

Alonso-Ramos P, Álvarez-Melo D, Strouhalova K, et al. (2021). The Cdc14 phosphatase controls resolution of recombination intermediates and crossover formation during meiosis. International Journal of Molecular Sciences 22, 9811. 10.3390/ijms22189811.

Attner MA, Amon A (2012). Control of the mitotic exit network during meiosis. Molecular Biology of the Cell 23, 3122–3132. 10.1091/mbc.E12-03-0235.

Briza P, Breitenbach M, Ellinger A, Segall J (1990a). Isolation of two developmentally regulated genes involved in spore wall maturation in Saccharomyces cerevisiae. Genes & Development 4, 1775–1789. 10.1101/gad.4.10.1775.

Briza P, Ellinger A, Winkler G, Breitenbach M (1990b). Characterization of a DL-dityrosine-containing macromolecule from yeast ascospore walls. Journal of Biological Chemistry 265, 15118–15123. 10.1016/S0021-9258(18)77231-1.

Briza P, Winkler G, Kalchhauser H, Breitenbach M (1986). Dityrosine is a prominent component of the yeast ascospore wall. Journal of Biological Chemistry 261, 4288–4294.

Broach JR, Strathern JN, Hicks JB (1979). Transformation in yeast: development of a hybrid cloning vector and isolation of the CAN1 gene. Gene 8, 121–133.

Buonomo SBC, Rabitsch KP, Fuchs J, et al. (2003). Division of the nucleolus and its release of Cdc14 during anaphase of meiosis I depends on separase, Spo12, and Slk19. Developmental Cell 4, 727–739.

Carrozza MJ, Li B, Florens L, et al. (2005). Histone H3 methylation by Set2 directs deacetylation of coding regions by Rpd3S to suppress spurious intragenic transcription. Cell 123, 581–592. 10.1016/j.cell.2005.10.023.

Chu S, DeRisi J, Eisen M, Mulholland J, Botstein D, Brown PO, Herskowitz I (1998). The transcriptional program of sporulation in budding yeast. Science 282, 699–705. 10.1126/science.282.5389.699.

Chu S, Herskowitz I (1998). Gametogenesis in yeast is regulated by a transcriptional cascade dependent on Ndt80. Molecular Cell 1, 685–696. 10.1016/S1097-2765(00)80068-4.

Culotti J, Hartwell LH (1971). Genetic control of the cell division cycle in yeast. III. Seven genes controlling nuclear division. Experimental Cell Research 67, 389–401. 10.1016/0014-4827(71)90424-1.

D’Amours D, Amon A (2004). At the interface between signaling and executing anaphase—Cdc14 and the FEAR network. Genes & Development 18, 2581–2595. 10.1101/gad.1247304.

Engel SR, Dietrich FS, Fisk DG, et al. (2014). The reference genome sequence of Saccharomyces cerevisiae: then and now. G3: Genes, Genomes, Genetics 4, 389–398. 10.1534/g3.113.008995.

Gietz RD, Schiestl RH (2007). Quick and easy yeast transformation using the LiAc/SS carrier DNA/PEG method. Nature Protocols 2, 35–37.

Holt LJ, Tuch BB, Villén J, Johnson AD, Gygi SP, Morgan DO (2009). Global analysis of Cdk1 substrate phosphorylation sites provides insights into evolution. Science 325, 1682–1686. 10.1126/science.1172867.

Hunter N (2015). Meiotic recombination: the essence of heredity. Cold Spring Harbor Perspectives in Biology 7, a016618. 10.1101/cshperspect.a016618.

Jaspersen SL, Charles JF, Tinker-Kulberg RL, Morgan DO (1998). A late mitotic regulatory network controlling cyclin destruction in Saccharomyces cerevisiae. Molecular Biology of the Cell 9, 2803–2817. 10.1091/mbc.9.10.2803.

Keeney S, Giroux CN, Kleckner N (1997). Meiosis-specific DNA double-strand breaks are catalyzed by Spo11, a member of a widely conserved protein family. Cell 88, 375–384. 10.1016/S0092-8674(00)81876-0.

Keogh M-C, Kurdistani SK, Morris SA, et al. (2005). Cotranscriptional Set2 methylation of histone H3 lysine 36 recruits a repressive Rpd3 complex. Cell 123, 593–605. 10.1016/j.cell.2005.10.025.

King GA, Goodman JS, Schick JG, et al. (2019). Meiotic cellular rejuvenation is coupled to nuclear remodeling in budding yeast. eLife 8, e47156. 10.7554/eLife.47156.

King GA, Wettstein R, Varberg JM, et al. (2023). Meiotic nuclear pore complex remodeling provides key insights into nuclear basket organization. Journal of Cell Biology 222, e202204039. 10.1083/jcb.202204039.

Koch BA, Staley E, Jin H, Yu H-G (2020). The ESCRT-III complex is required for nuclear pore complex sequestration and regulates gamete replicative lifespan in budding yeast meiosis. Nucleus 11, 219–236. 10.1080/19491034.2020.1812872.

Lam I, Keeney S (2015). Mechanism and regulation of meiotic recombination initiation. Cold Spring Harbor Perspectives in Biology 7, a016634. 10.1101/cshperspect.a016634.

Li B, Gogol M, Carey M, Lee D, Seidel C, Workman JL (2007). Combined action of PHD and chromo domains directs the Rpd3S HDAC to transcribed chromatin. Science 316, 1050–1054. 10.1126/science.1139004.

Marston AL, Amon A (2004). Meiosis: cell-cycle controls shuffle and deal. Nature Reviews Molecular Cell Biology 5, 983–997. 10.1038/nrm1526.

Marston AL, Lee BH, Amon A (2003). The Cdc14 phosphatase and the FEAR network control meiotic spindle disassembly and chromosome segregation. Developmental Cell 4, 711–726.

Martín H, Arroyo J, Sánchez M, Molina M, Nombela C (1993). Activity of the yeast MAP kinase homologue Slt2 is critically required for cell integrity at 37 degrees C. Molecular and General Genetics 241, 177–184. 10.1007/BF00280215.

Matos J, Blanco MG, Maslen S, Skehel JM, West SC (2011). Regulatory control of the resolution of DNA recombination intermediates during meiosis and mitosis. Cell 147, 158–172. 10.1016/j.cell.2011.08.032.

Miao M, Ryan KJ, Wente SR (2006). The integral membrane protein Pom34p functionally links nucleoporin subcomplexes. Genetics 172, 1441–1457. 10.1534/genetics.105.052068.

Nasmyth KA, Tatchell K (1980). The structure of transposable yeast mating type loci. Cell 19, 753–764.

Neiman AM (2011). Sporulation in the budding yeast Saccharomyces cerevisiae. Genetics 189, 737–765.

Onischenko E, Stanton LH, Madrid AS, Kieselbach T, Weis K (2009). Role of the Ndc1 interaction network in yeast nuclear pore complex assembly and maintenance. Journal of Cell Biology 185, 475–491. 10.1083/jcb.200810030.

Pablo-Hernando ME, Arnaiz-Pita Y, Nakanishi H, et al. (2007). Cdc15 is required for spore morphogenesis independently of Cdc14 in Saccharomyces cerevisiae. Genetics 177, 281–293. 10.1534/genetics.107.076133.

Rahal R, Amon A (2008). The Polo-like kinase Cdc5 interacts with FEAR network components and Cdc14. Cell Cycle 7, 3262–3272. 10.4161/cc.7.20.6852.

Rock JM, Amon A (2009). The FEAR network. Current Biology 19, R1063–R1068. 10.1016/j.cub.2009.10.002.

Rodríguez-Jiménez P, Alonso-Ramos P, Acosta I, Álvarez-Melo D, Carballo JA (2026). Nucleolar Cdc14 splitting reflects recombination context and meiotic chromosome dynamics. International Journal of Molecular Sciences 27, 888. 10.3390/ijms27020888.

Ruediger CT, Styler BS, Sawyer EM, et al. (2026). Tom70-mediated mitochondria–nuclear envelope contacts regulate nuclear pore complex inheritance during gametogenesis. Nature Communications (published online 23 October 2026). 10.1038/s41467-026-78562-1.

Schindelin J, Arganda-Carreras I, Frise E, et al. (2012). Fiji: an open-source platform for biological-image analysis. Nature Methods 9, 676–682. 10.1038/nmeth.2019.

Schneider CA, Rasband WS, Eliceiri KW (2012). NIH Image to ImageJ: 25 years of image analysis. Nature Methods 9, 671–675. 10.1038/nmeth.2089.

Shou W, Azzam R, Chen SL, et al. (2002). Cdc5 influences phosphorylation of Net1 and disassembly of the RENT complex. BMC Molecular Biology 3, 3. 10.1186/1471-2199-3-3.

Shou W, Seol JH, Shevchenko A, et al. (1999). Exit from mitosis is triggered by Tem1-dependent release of the protein phosphatase Cdc14 from nucleolar RENT complex. Cell 97, 233–244. 10.1016/S0092-8674(00)80733-3.

Stegmeier F, Amon A (2004). Closing mitosis: the functions of the Cdc14 phosphatase and its regulation. Annual Review of Genetics 38, 203–232. 10.1146/annurev.genet.38.072902.093051.

Visintin C, Tomson BN, Rahal R, et al. (2008). APC/C-Cdh1-mediated degradation of the Polo kinase Cdc5 promotes the return of Cdc14 into the nucleolus. Genes & Development 22, 79–90. 10.1101/gad.1601308.

Visintin R, Hwang ES, Amon A (1999). Cfi1 prevents premature exit from mitosis by anchoring Cdc14 phosphatase in the nucleolus. Nature 398, 818–823. 10.1038/19775.

Yeheskely-Hayon D, Kotler A, Stark M, Hashimshony T, Sagee S, Kassir Y (2013). The roles of the catalytic and noncatalytic activities of Rpd3L and Rpd3S in the regulation of gene transcription in yeast. PLOS ONE 8, e85088. 10.1371/journal.pone.0085088.

Zickler D, Kleckner N (1999). Meiotic chromosomes: integrating structure and function. Annual Review of Genetics 33, 603–754. 10.1146/annurev.genet.33.1.603.

