## Supplemental Data 1 for "UV-excited autofluorescence uncovers multicopy suppressors of a Cdc14-associated meiotic defect"

### Figure S1

**A**

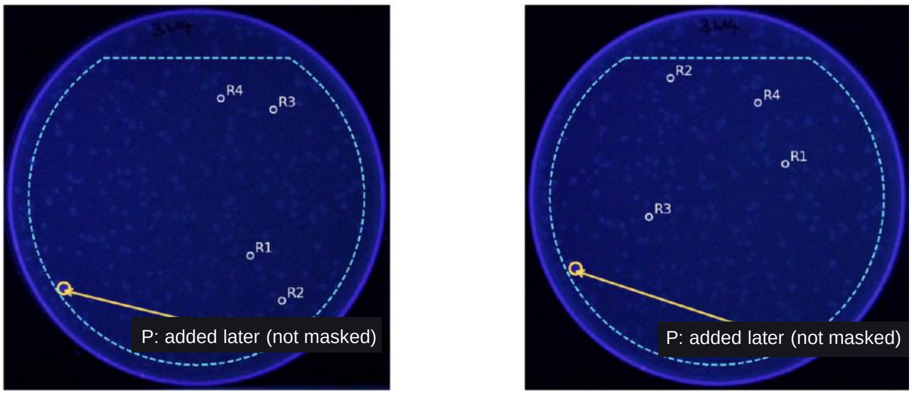


**Peripheral-feature omission and local correction**

P: (1270,1040) in IMG_4943(2).JPG; (506,974) in IMG_4946(1).JPG (2048-px originals)

Original 90% mask retained; P required a separate 95% local geometric guard. No retrospective automatic detection. P was added to the later local evaluation; it was absent from the original automatic mask.

**B**

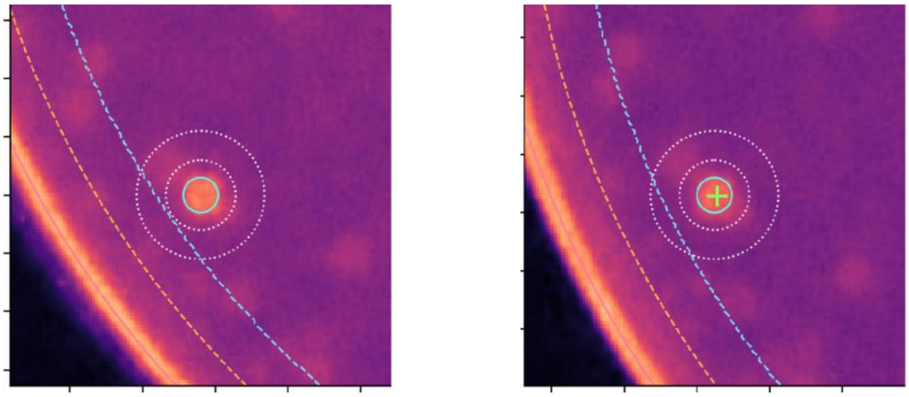

**Pseudocolour, ROI aperture and perimeter geometry**

6-px core · 12–22-px annulus · 90% original mask · local 95% rim guard; original blue (8-bit JPEG DN).

**C**

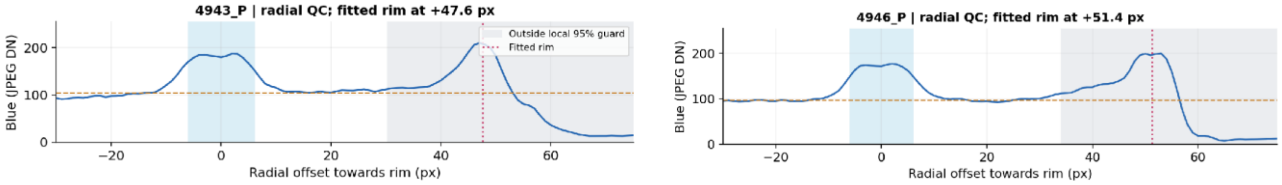


**Radial blue-intensity profiles distinguishing punctum and rim (W07 P2)**

Fitted rim is diagnostic only; the local background annulus excludes it. Samples are not biological replicates.

**Supplementary Figure S1 | Local fluorescence measurements and assessment of peripheral signals.** (A) Blue-channel views of the same query plate in two acquisitions, showing the peripheral fluorescence-associated feature P and its position relative to the original detection boundary. P was not selected by the original 90% automatic plate mask and was assessed separately using a local geometric extension. (B) Pseudocolour representations showing the local signal region, surrounding background annulus and plate-perimeter geometry. The analysis used a 6-pixel signal radius and a background annulus extending from 12 to 22 pixels. (C) Radial blue-intensity profiles from the two acquisitions, illustrating the distinction between the localised feature and the bright plate rim. The fitted rim was used for diagnostic assessment and excluded from the local background measurement. P was not assigned to a historically identified or genetically validated colony.

### Figure S2

A

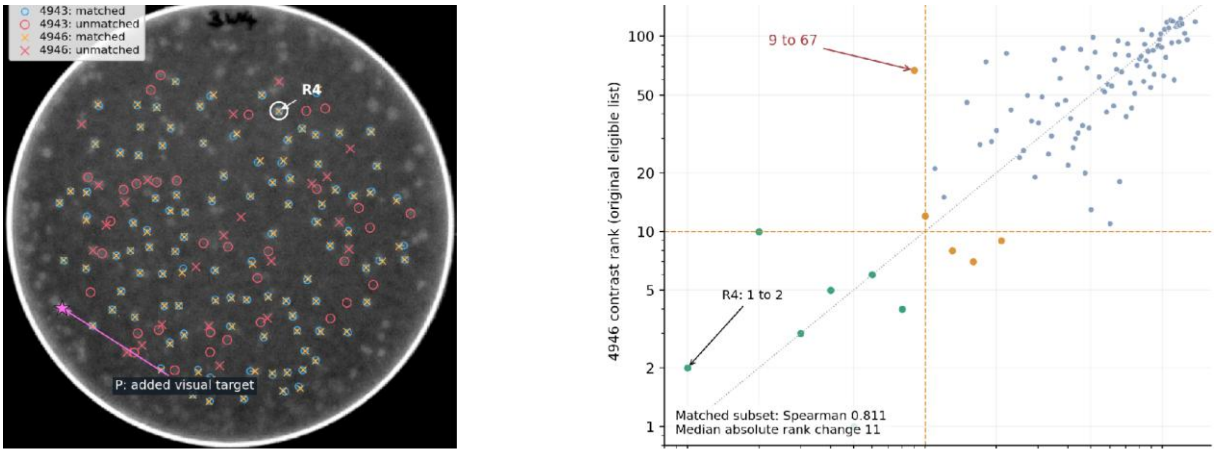


| Scope | Counts 4943 / 4946 | Matched pairs | Unmatched 4943 / 4946 | 4943 top 10 found in 4946 | 4946 top 10 found in 4943 | Strict top-10 intersection |
|---|---|---|---|---|---|---|
| Original eligibility | 137 / 127 | 104 | 33 / 23 | 9/10 | 10/10 | 7/10 |
| Including added P | 138 / 128 | 105 | 33 / 23 | 9/10 | 10/10 | 7/10 |

B

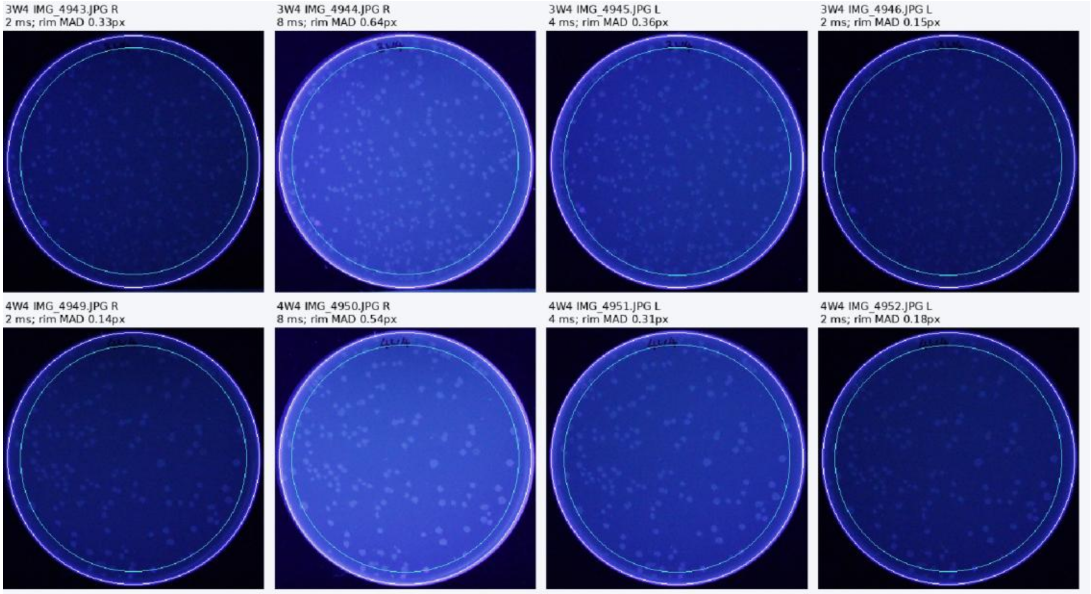


C

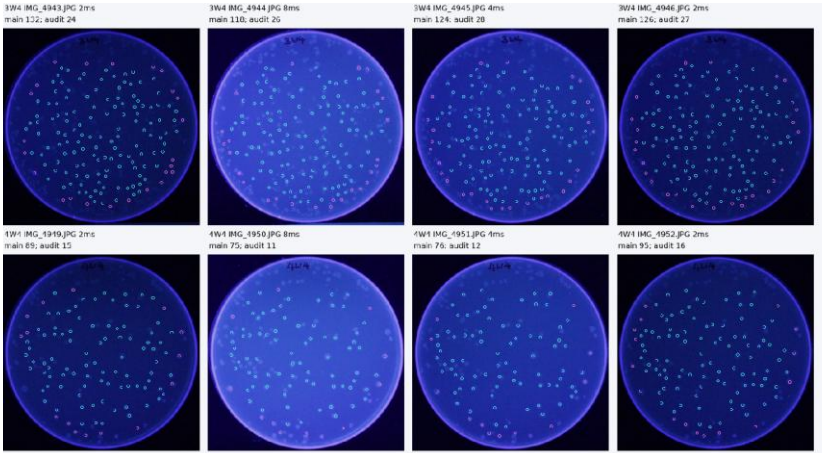

**Supplementary Figure S2 | Detection correspondence and acquisition quality control across UV screening photographs.**(A) Expanded spatial-correspondence analysis of two photographs from the 3W4 development plate. The original detection lists contained 137 and 127 maxima, with 104 spatial matches and seven shared top-ten detections. Separate inclusion of the locally assessed peripheral feature P increased the lists to 138 and 128 maxima and the matched count to 105, without changing the top-ten overlap. (B) Representative plate-geometry and image-registration checks for four acquisitions each of the 3W4 development plate and 4W4 comparison plate. (C) Corresponding detection and peripheral-feature quality-control examples. The peripheral audit identified out-of-mask features with contrast exceeding the main-mask maximum in 9 of 24 additional photographs. The main cross-plate detection measurements were retained unchanged. Panels B and C show representative acquisitions from two plate families, not the complete seven-family dataset. Repeated photographs and their pairwise comparisons are technical observations, not independent biological replicates.

### Figure S3

A

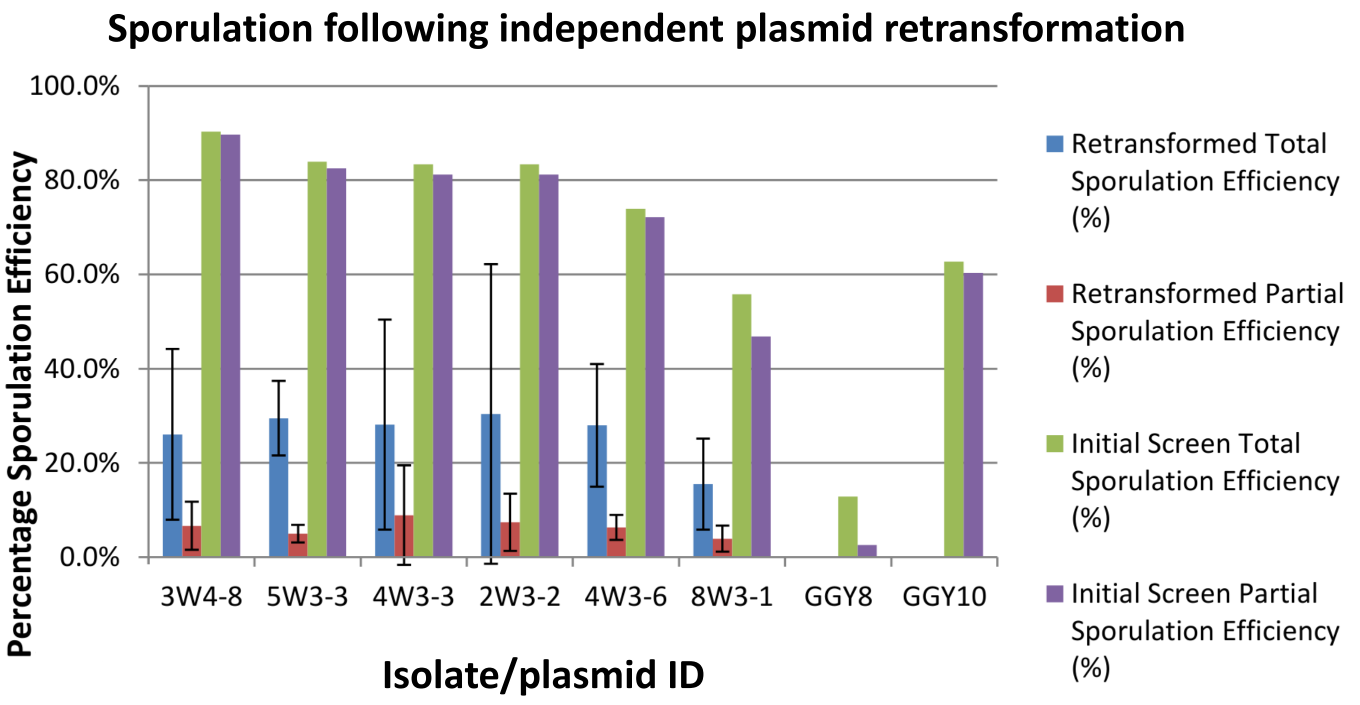


B

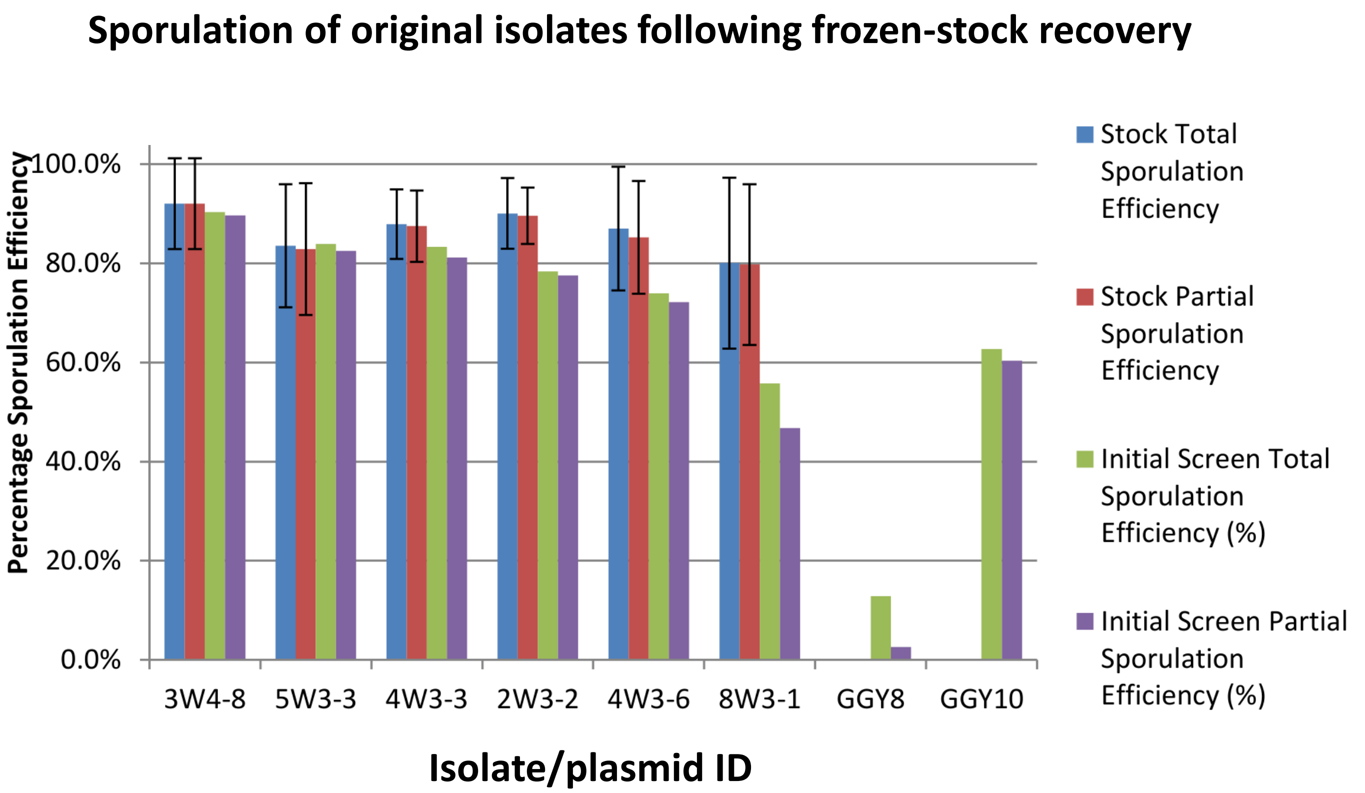


**Supplementary Figure S3 | Sporulation phenotypes of six primary-screen isolates and their recovered plasmids.** (A) Sporulation efficiencies of six isolates selected from the primary screen, compared with those obtained following independent retransformation of their recovered plasmids into fresh GGY8 recipients. None of the retransformed plasmids reproduced the elevated sporulation observed in the original isolates. Untransformed GGY8 and GGY10 controls are included.(B) Sporulation efficiencies of the same six isolates following recovery from frozen stocks, compared with their initial screening measurements. The original isolates retained elevated sporulation, in contrast to the lack of suppression observed upon plasmid retransformation. Error bars represent ±2 standard deviations (SD). Repeated measurements do not represent additional independent transformants.

#### Supplementary Table S1A | Yeast strains used in the screen

| Strain | Background / ploidy | Genotype | Use in this study |
|---|---|---|---|
| GGY8 | SK1 · diploid | *MATa/MATα*, *ura3*, *ho*::*LYS2/"*, *lys2/"*, *leu2*::*hisG/"*, *his4x/"*, *CDC14*-3HA-KanMX4/" | Homozygous tagged *cdc14-HA3* query; library transformation and plasmid retransformation |
| GGY10 | SK1 · diploid | *MATa/MATα*, *ura3/"*, *ho*::*LYS2/"*, *lys2/"*, *leu2*::*hisG/"*, *his4x/"*, *CDC14/CDC14*-3HA-KanMX4 | Heterozygous *CDC14/cdc14-HA3* reference; UV-plate and patch controls |

SK1-derived diploids used for primary library screening and independent plasmid assays.

#### Supplementary Table S1B | Plasmids and mapped genomic-library inserts

| PLASMID / ISOLATE | ORIGINAL VECTOR | GENETIC CONTENT / MAPPED INTERVAL | SOURCE / ARCHIVED MAP |
|---|---|---|---|
| YEP13 (EMPTY VECTOR) | YEp13; 2µ, LEU2 | Unmodified multicopy library backbone; empty-vector reference | screen records |
| YEP13 GENOMIC LIBRARY | YEp13; 2µ, LEU2 | Partially digested AB320 S. cerevisiae genomic DNA cloned at BamHI | Broach et al., 1979 |
| PD.2 | pRS426; 2µ, URA3 | *CDC14*–HA3 high-copy reference construct | Alonso-Ramos et al., 2021 |
| YEP352-*SLT2* | YEp352; 2µ, URA3 | *SLT2* and flanking sequences; reported ~2.2-kb SalI–EcoRI fragment | Humberto Martín (gift); Methods |
| 5W4-5 *POM34* | YEp13, via 5W4-5 | *POM34* intact; predicted *BRE2–MEU1* fusion after NcoI deletion/religation | p5W4-5_POM34(2).xdna; screen records |
| 3W4-8 | YEp13; 2µ, LEU2 | chrXIV:335,352–344,598; *YGP1*, *PGA1*, *IGO1*, YNL155W, *NSG2*, *ASI2*, Truncated *YCK2* | xDNA 3W4-8 |
| 5W3-3 | YEp13; 2µ, LEU2 | chrV:450,011–454,512; *RTR1*, *EMP65*, and truncated *COX15* | xDNA 5W3-3 |
| 4W3-3 | YEp13; 2µ, LEU2 | chrXVI:358,161–362,551; A - *RED1*, *TMA7 YPT6*, dubious *VPS63*; B - Truncated *FMP30*, dubious YPL102C, *ELP4*, truncated *ATG21* | xDNA 4W3-3A/4W3-3B* |
| 2W3-2 | YEp13; 2µ, LEU2 | chrIX:299,472–304,430; *EMC5*, Dubious YIL028W, Dubious YIL029W-A, tK(CUU)I, YILCdelta3, YIL029C, Truncated *SSM4* | xDNA 2W3-2† |
| 4W3-6 | YEp13; 2µ, LEU2 | chrX:682,073–686,478; Truncated *MET5*, truncated *IML1*, ARS1021 also present | xDNA 4W3-6 |
| 8W3-1 | YEp13; 2µ, LEU2 | chrV:450,011–454,512; *RTR1*, *EMP65*, and truncated *COX15* | xDNA 8W3-1 |
| 10W4-11 | YEp13; 2µ, LEU2 | chrXV:913,464–920,498; *PMT3*, *GNT1*, Truncated *LDB19* | xDNA 10W4-11C* |
| 5W4-5 | YEp13; 2µ, LEU2 | chrXII:174,847–180,279; *POM34*, *PML1*, *BRE2*, *MEU1* (intact); *PPR1* (truncated) | xDNA 5W4-5 |
| 10W4-17 | YEp13; 2µ, LEU2 | chrVIII:164,487–170,510; *DAP2*, *YHI9*, *SLT2* (intact); RPN1 (truncated); YHR028W-A (dubious); ARS809 | xDNA 10W4-17 |
| 6W4-8 | YEp13; 2µ, LEU2 | chrXIII:413,819–418,734; *RCO1* (intact); *PDS5* (truncated); YMR075C-A (dubious) | xDNA 6W4-8 |
| 8W3-16 | YEp13; 2µ, LEU2 | chrXV:1,011,631–1,020,719; Truncated *VTS1*, *PDE2*, *PRT1*, *PRE10*, truncated *PIP2* | xDNA 8W3-16A* |
| 9W4-4 | YEp13; 2µ, LEU2 | chrXIII:477,651–484,718; YMR105W-A, *SPG4*, Truncated ARS1317, YMRWdelta15, YMRWdelta14, Truncated ILV2, *YKU80*, tK(CUU)M | xDNA 9W4-4D* |
| 11W4-9 | YEp13; 2µ, LEU2 | chrXII:708,105–718,268; *GUF1*, Dubious YLR286W-A, YLR287C, *RPS30A*, *MEC3*, *CTS1*, YLR290C, YLR285C-A | xDNA 11W4-9 |
| 12W3-16 | YEp13; 2µ, LEU2 | chrXIV:363,328–365,354; Truncated THO2 | xDNA 12W3-16 |
| 9W3-4 | YEp13; 2µ, LEU2 | chrIV:391,787–396,572; *SLM3*, dubious YDL032W, Truncated *GPR1*, Truncated *DBP10*, Truncated dubious YDL034W | xDNA 9W3-4/9W3-4A* |
| 2W1-15 | YEp13; 2µ, LEU2 | chrX:103,711–109,986; *QCR8*, *HAL5*, *ERG20*, Truncated *SET2*, Truncated *TRK1* | xDNA 2W1-15 |
| 9W4-2 | YEp13; 2µ, LEU2 | chrXV:538,525–545,086; *RPO31*, *TRS33*, Truncated *RPT5* | xDNA 9W4-2 |
| 2W1-1 | YEp13; 2µ, LEU2 | chrVII:715,575–723,110; *SHY1*, *DAM1*, uncharacterized YGR111W, Truncated *SPT6*, dubious YGR115C &YGR114C | xDNA 2W1-1 |
| 4W1-15 | YEp13; 2µ, LEU2 | chrVII:968,363–970,608; *PEX21*, Truncated *KEL2*, Truncated *PFK1* | xDNA 4W1-15 |
| 10W4-18 | YEp13; 2µ, LEU2 | chrVI:78,706–83,877; *CAF16*, *GYP8*, *STE2*, Truncated *CAK1* | xDNA 10W4-18 |

**Notes.** Genomic coordinates and annotations follow the archived screen maps. Insert interiors were inferred from reference-genome mapping of terminal Sanger reads; complete insert sequences were not experimentally read end-to-end.

***** Suffixes A/B/C/D denote archived map labels; equivalence with an unsuffixed historical isolate or a particular retransformed plasmid is not established by the filename alone. **†** The 2W3-2 archived MboI map filename also contains a GATC label "2W3-3".

**Supplementary Table S2 | Primary-screen candidate records**

The 20 highest-ranked corrected GGY8 query records by partial sporulation. Primary measurements are shown independently of later plasmid validation.

| Rank | Primary isolate | Total sporulation (%) | Partial sporulation (%) | G418 segregation | Independent plasmid follow-up |
|---|---|---|---|---|---|
| **1** | 3W4-8 | 90.32 | 89.68 | 2:2 | High primary phenotype not reproduced |
| **2** | 5W3-3 | 83.94 | 82.48 | 2:2 | High primary phenotype not reproduced |
| **3** | 4W3-3 | 83.33 | 81.16 | 2:2 | High primary phenotype not reproduced |
| **4** | 2W3-2 | 78.40 | 77.60 | 2:2 | High primary phenotype not reproduced |
| **5** | 4W3-6 | 73.91 | 72.17 | 2:2 | High primary phenotype not reproduced |
| **6** | 8W3-1 | 55.77 | 46.79 | 2:2 | High primary phenotype not reproduced |
| **7** | 10W4-11 | 41.23 | 39.47 | 4:0 | Not confirmed after adjustment |
| **8** | 5W4-5 | 42.20 | 33.94 | 4:0 | Confirmed in fresh GGY8 |
| **9** | 10W4-17 | 40.98 | 32.79 | 4:0 | Confirmed in fresh GGY8 |
| **10** | 6W4-8 | 42.86 | 29.41 | 4:0 | Confirmed in fresh GGY8 |
| **11** | 8W3-16 | 44.36 | 28.57 | 4:0 | Not assayed independently |
| **12** | 9W4-4 | 51.56 | 28.12 | 4:0 | Quantitative assay: 9W4-4D |
| **13** | 11W4-9 | 43.94 | 25.00 | 4:0 | Not assayed independently |
| **14** | 12W3-16 | 51.38 | 23.76 | 4:0 | Not assayed independently |
| **15** | 9W3-4 | 44.83 | 22.07 | Not documented | Not assayed independently |
| **16** | 2W1-15 | 42.40 | 20.80 | 4:0 | Not assayed independently |
| **17** | 9W4-2 | 33.33 | 20.74 | 4:0 | Not assayed independently |
| **18** | 2W1-1 | 40.91 | 20.45 | 4:0 | Not assayed independently |
| **19** | 4W1-15 | 30.71 | 20.00 | 4:0 | Not assayed independently |
| **20** | 10W4-18 | 31.82 | 19.70 | 4:0 | Not assayed independently |

Primary values: original Complete Sporulation Efficiencies; total sporulation includes monads, while partial sporulation excludes them. Values are percentages rounded to two decimals. G418 segregation: six initially high-sporulation isolates showed 2:2. Thirteen additional top-20 records have documented 4:0 results.
